# Back to the future: a refined single user photostation for massively scaling herbarium digitization

**DOI:** 10.1101/2020.09.09.288506

**Authors:** Charles C. Davis, Jonathan A. Kennedy, Christopher J. Grassa

## Abstract

The digitization and online mobilization of herbarium specimens has greatly facilitated their access and helped ignite a revolution in the biodiversity sciences (Drew et al., 2017; Hedrick et al., 2020; Nelson et al., 2015; Soltis, 2017; Sweeney et al., 2018; Thiers et al., 2016). These efforts have mobilized millions of specimens with significant economies of scale and accelerated advances in scientific investigations, including phenological studies of climate change, species range assessments, and biotic interactions (Hedrick et al., 2020; Meineke et al., 2019; Meineke et al., 2018; Pearson et al., 2020; Willis et al., 2017). In addition, the use of natural history collections to answer scientific questions using only their digitized representation, rather than the physical specimen itself–i.e., Digitization 2.0 *sensu* Hedrick et al. (2020)–has sparked the integration and development of new scholarly disciplines and lines of inquiry not previously possible. Despite these exciting new directions, however, Digitization 1.0 *sensu* Hedrick et al. (2020)–i.e., the generation of digitized products from the physical specimen–remains an active area of innovation and development. This relates to both hardware and workflow innovations as well as their integration with advancements in software. Along these lines, innovations in these areas have greatly increased the cost-effectiveness of digitizing herbarium specimens and enabled the successful mobilization of entire collections and whole floristic regions (Heerlien et al., 2015; Pignal and Michiels, 2012; Schorn et al., 2016; Slijkhuis, 2014; Sweeney et al., 2018; van Oever and Gofferjé, 2012). Here, we present a novel photostation and workstation design for imaging herbarium specimen that represents a dramatic improvement upon existing approaches and is scalable for large and small institutions alike.

## Conveyor belt technologies and herbarium digitization

Herbarium digitization has historically been accomplished using light boxes, light tables, or scanning setups, which are operated from a single user interface (see overview of traditional approaches by Nelson et al., 2015). A major breakthrough in the digitization of herbaria was the application of conveyor belt technologies (Heerlien et al., 2015; Pignal and Michiels, 2012; Schorn et al., 2016; Slijkhuis, 2014; Sweeney et al., 2018; van Oever and Gofferjé, 2012). Such conveyor belt setups are industrial-scale, high-throughput systems dedicated especially to vascular plant herbarium specimens mounted to sheets. A central feature of these systems is an automated conveyor belt apparatus, which allows image capture and specimen handling to occur simultaneously, and which prioritizes imaging and the capture of at least minimal specimen data. To date, these efforts have been successfully deployed at a massive scale at the Muséum national d’Histoire naturelle Paris (P) (Le Bras et al., 2017; Slijkhuis, 2014), the Naturalis Biodiveristy Center (NL) (Rogers, 2016; Slijkhuis, 2014), the United States National Herbarium (US), and at our home institution, the Harvard University Herbaria (HUH) (Schorn et al., 2016; Sweeney et al., 2018). It is also worth mentioning that the vendor Picturae [https://picturae.com/en/what-we-offer/digitizing-collections], who offers solutions using a conveyor-belt workflow (Slijkhuis, 2014), was contracted for the successful digitization of P (Le Bras et al., 2017), NL (Heerlien et al., 2015), and more recently US.

The HUH has played a role in innovating rapid, efficient imaging and transcription workflows (Schorn et al., 2016; Sweeney et al., 2018). In particular, as part of the ‘Mobilizing New England Vascular Plant Specimen Data to Track Environmental Changes Project’ (NEVP), a Thematic Collections Network (TCN) funded through the U.S. National Science Foundation’s Advancing the Digitization of Biodiversity Collections (ADBC) program (award 1208835), the HUH (in collaboration with colleagues at Yale University [Patrick Sweeney] and the University of South Carolina [Bili Starly]) pioneered a novel and cost-effective conveyor belt system to digitize (image, transcribe, and georeference) the entirety of the HUH’s ∼350,000 New England vascular plant holdings (Schorn et al., 2016; Sweeney et al., 2018). This was a dramatic improvement from previous efforts in the herbarium community, which utilized scanning setups, photo tables, and light boxes (Nelson et al., 2015). At the same time, these conveyor belts were expensive and required a large space footprint. In deploying the conveyor belts at the HUH, the physical space requirements meant the digitization area was not immediately co-located with the collection. Distance between the collection and the digitization area was a barrier to reducing transit time and also increased the risk of specimen damage during transit. In the case of the HUH, the digitization area was located in the same building and in a stringent pest-management zone. However, had the available space necessitated locating the digitization area further from the collection, this would have introduced the need for additional pest-control procedures adding significant time and labor to the workflow. Moreover, each conveyor belt relied on the coordinated effort of minimally two staff members and absences caused a measurable decline in efficiency. With further analysis, our team discovered that many of the positive efficiencies of the conveyor belt system were the result of ergonomic and workflow-specific customizations which were not specific to the use of a conveyor belt system. Taking this entire workflow into consideration, we noted that the presence of a conveyor belt did not reduce the necessary physical motions for an operator to stage and de-stage a specimen for imaging compared to other single-user workflows, nor did it increase the speed of that activity. Our team hypothesized that a single-user photostation, better optimized for herbarium imaging workflows, would produce similar or greater efficiencies while resolving the space and coordination challenges of the conveyor belt system. At the same time, the team conceived of leveraging new software tools in the digitization process, so that the imaging workflow could be designed and optimized independent of additional data capture-related activities.

## Returning to a single-operator photostation for rapid herbarium specimen imaging

With continued investment in process and workflow analysis, we developed a single-operator photostation specifically designed and fabricated for herbarium specimen digitization (Fig. 1). This photostation design has now been in operation at the HUH since mid-2018 (first prototype deployed fall of 2017) and has greatly exceeded expectations for increased imaging efficiencies while also realizing the benefits of a small, flexible, and scalable system. In a nutshell, we have taken herbarium digitization efforts ‘Back to the future’, returning to a more traditional single user approach, but with much greater efficiency that is scalable to small and large collections alike. Here, we present our workflow and novel photostation design in the hopes that similar institutions, large and small, can benefit from our efforts.

**Figure 1.**
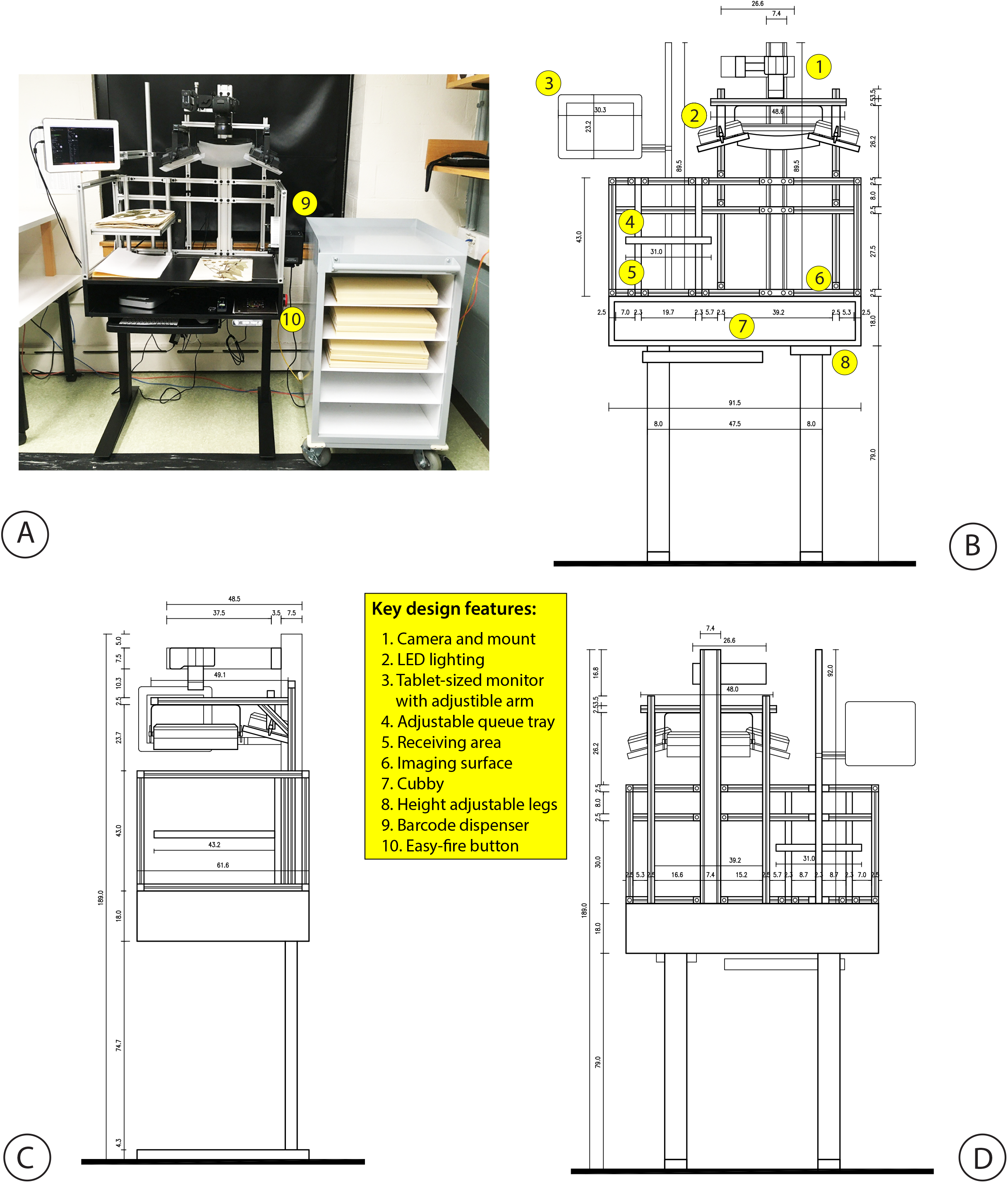
Single-user photostation design. 1) Photograph of photostation. Technical drawings of front (2), side (3), and (4) rear view of station. Measurements in metric units (cm). Key details highlighted in yellow (2) with numbered legend. Built-in jig for mounting specimens with ruler and color checker, automated barcode dispenser, and easy-fire button shown only in photograph on imaging table.

Our photostation design presented in Figure 1 can be constructed using available consumer hardware. The upper frame uses Grainger extruded aluminum framing and fixtures, while the cubby is made from laminated wood. Automated height-adjustable Fully Jarvis standing desk legs are attached to the base of the photostation. Multiple camera technologies can be attached to the photostation and easily upgraded over time. The HUH presently utilizes the Canon 5DSR camera body which is capable of 50MP images, equivalent to a 520ppi image of a standard herbarium sheet, and ZEISS Milvus 50mm f/2 macro lens. Genaray 5600K LED lighting with a >90 CRI rating and a controlling computer (Mac Mini) with small tablet-sized monitor (mounted at eye level) are included in the hardware configuration. Capture One software is used for image capture on the photostation as well as post-processing on a separate computer. Construction of the photostation requires tools for cutting wood and extruded aluminum framing. Drilling is necessary to attach the frame and legs to the wooden cubby. Assembly of the upper frame and attachment of the electronic components is accomplished with Grainger fasteners and an Allen wrench. Institutions without local expertise can work with local builders or hackerspaces for the construction. After the completion and production testing of our prototype, the HUH contracted a local builder to fabricate six photostations using our final design. The total cost of the materials, electronic components, and software is under $10,000, while the cost of labor will vary by region.

## Photostation workflow

Alongside our novel photostation hardware design, we also greatly optimized our workflow (Fig. 2). This imaging workflow takes place as follows: 1) A batch of specimens are brought to the digitization area in a manner appropriate for the specific collection and distance. In the case of the HUH, specimen folders are removed from cabinets and loaded into an herbarium cart that is transported a short distance to the digitization area, generally to a room on the same floor. Photostations are deployed on multiple floors to reduce transit time. 2) Before imaging, specimens are checked for damage, minor repairs are performed as needed, and specimens with severe damage are set aside. Any specimen missing a barcode has one affixed. As the specimens are inspected, they are stacked so that their order is reversed prior to imaging. 3) An imaging session begins by logging into the computer attached to the photostation, launching the photo capture software and creating a new session named for the operator and date. Once the attached camera and lights are turned on, a level is placed underneath the camera lens to check the tilt along the horizontal and z-axis, the camera lens is focused using a focus chart on the imaging stage, the camera settings are checked (shutter speed, ISO, aperture), and a test image is captured using a large color checker (X-Rite ColorChecker Digital SG). Wireless mouse and keyboard, level, focus chart, and color checker are stored in a cubby directly below the photo stage for quick access. 4) To begin imaging, a specimen folder is placed on a “queue” shelf directly next to the imaging stage. 5) The specimens are removed from the folder and kept on the queue shelf while the folder is placed on the imaging stage. 6) An image is captured using an easy-fire button adjacent to the stage, attached to the outside of the photostation frame. This placement helps ensure the hand of the operator is out of the camera’s view. 7) After hearing the audible shutter-sound (screen flash can also be used for the hearing impaired), the operator moves the folder to the area immediately below the queue shelf (the receiving area). 8) The waiting stack of specimens are similarly imaged one by one, in rapid succession, as are any species covers. Images of the folder, species covers, and specimens will be preserved in sequence for later transcription. 9) A specimen is taken from the top of the specimen stack on the queue shelf and placed on the photo stage. A built-in jig, sized for standard 16.5” x 11.5” herbarium sheets, helps to quickly guide the placement of the specimen sheet. The HUH logo and mini color checkers are already attached to the jig to eliminate additional placements by the operator. 10) If a specimen did not receive a barcode ahead of imaging, a barcode is retrieved from an automatic barcode dispenser attached directly to the right of the photo stage. 11) After an image is captured, the operator can inspect the image using the monitor attached to the photostation. This tablet-sized monitor is only intended to identify gross problems with the image, such as camera skew or color distortion, that might be caused by an equipment failure or setup mistake. Because image capture can occur many times faster than the images can be rendered to the monitor, operators are only asked to inspect images at the start of the session and only occasionally thereafter. As specimens are imaged, they are stacked in the receiving area to the left of the photostage and under the queue tray. This stacking also corrects the ordering that was reversed during the curation step.

**Figure 2.**
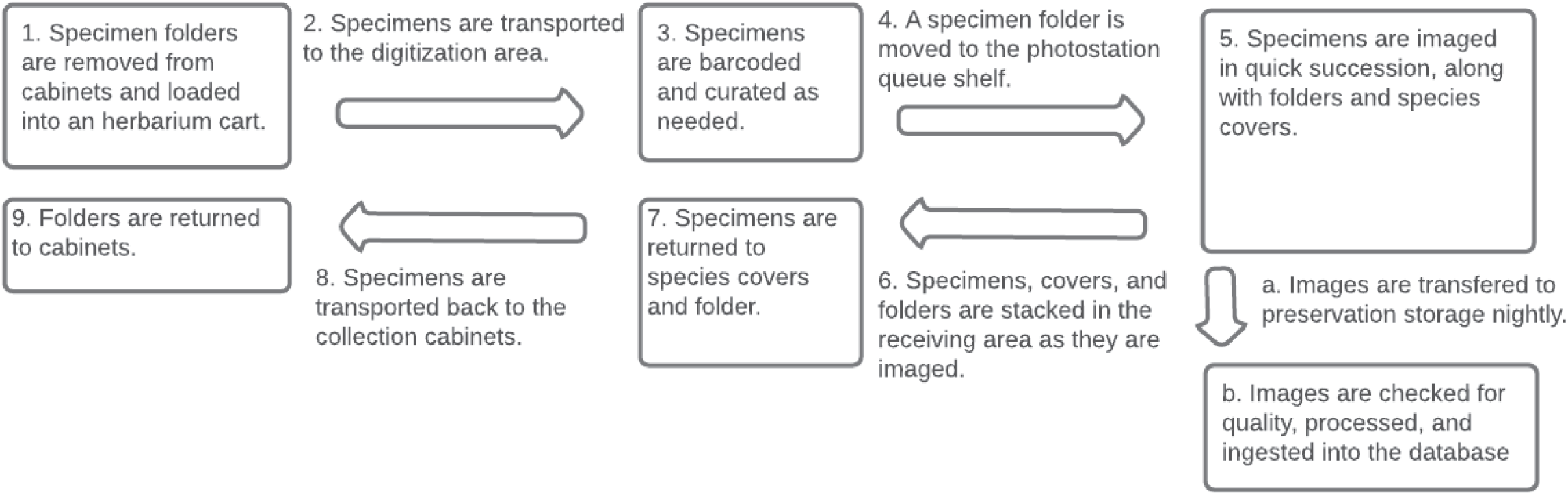
HUH digitization workflow.

The imaging workflow described above occurs very rapidly. After the initial setup procedure and inspection of the first image, capture of subsequent images can occur in as quickly as 4 seconds per image, although careful handling of fragile specimens and the number of species covers will raise the average rate of imaging. We have also observed that taxonomic identity can be a factor in the rate of image production, with thin, densely stacked specimens (e.g., grasses and orchids) more quickly imaged than thicker, woody specimens (e.g., many rosid tree species). The photostation design can also accommodate bulky specimen types (e.g., fruits and cacti), assuming the specimen fits onto the photostage, by adjusting the camera settings to create a large depth of field. Imaging sessions can include any number of images, though from our experience batches of fewer than 1,000 images are easier to review, quality control, post-process, and synchronize across networks and storage devices. Creating a new session is quickly accomplished within the image capture software and can be done at any point between specimen folders to limit the size of each batch.

## Ergonomic enhancements

Ergonomics were considered extensively during the design of the photostation, especially because the workflow calls for significant time performing highly repetitive motions (specimen placement, button press, specimen removal). To increase comfort, the photostation is built with adjustable height legs controlled with an electronic panel preset for up to seven users. Operators can stand on an ergonomic mat, or use a chair or stool of their choice, or alternate between sitting and standing. Significant consideration was also given to lighting. Many herbaria use light boxes or light tables for their imaging workflows. Lightboxes allow for controlled lighting of the photostage without interference from environmental light sources, such as windows and overhead lighting. In contrast, the use of light tables often necessitates a dark room. Because the quickness of staging and de-staging the specimen is a central factor for imaging efficiency, our photostation uses an open design and is subject to room lighting conditions. Our initial prototype required the room to be dark and a fabric hood was placed over the top of the photostation to reduce the brightness experienced by the operator. Operators were also supplied with optional OSHA-compliant partially tinted glasses to further reduce eyestrain. This worked but was less than ideal for operator comfort. We ultimately abandoned the dark room and installed overhead lighting that reasonably matched the color temperature of the lighting attached to the photostation. Because both sources of lighting evenly illuminate the stage, we have observed no problems with achieving a consistent white balance for accurate color reproduction. This approach significantly improved comfort and overall operator satisfaction while avoiding the inefficiencies of repeatedly opening, closing, and navigating around a light-box. We conducted numerous tests with the help of HUH curatorial staff throughout the development of our first two prototypes and final design. We concluded that 2-3 hours was the upper limit for comfortably imaging in a single session. Imaging at or above two hours, more than once per day, was considered to be both physically and mentally taxing. Fortunately, owing to the overall efficiency of the system, HUH staff regularly perform only one or two imaging sessions per week. This produces a sufficient number of images so that the remaining work-week can be dedicated to transcribing those images.

## Photostation production rates

Images produced by the photostation are timestamped, creating a lasting record of the system’s imaging rates. By comparing the timestamps of the first and last images in a session, it is trivial to calculate the average seconds per image for that session. To determine our real-world efficiencies, we analysed the time data from batches spanning different operators and taxonomic groups and were able to draw strong conclusions about the average efficiency of the imaging workflow. Using data from eight full-time digitizers and five full-time staff who digitize part-time, spanning approximately two years (corresponding to the deployment of the final iteration of the photostation design), we calculated an average imaging rate of 8 seconds per image. This rate does not include photostation setup, which is performed by the operator in about five minutes. The above rate also does not include time spent transporting, curating, and refiling specimens, and does include images taken of folders and species covers. Many of these factors are highly variable between institution and we believe our timing analysis is comparable to those published for other imaging workflows.

The speed of imaging is throttled by the physical speed of the operator, therefore decisions regarding personnel, work organization, and staff management can make significant impacts on achieving higher efficiencies. However, the HUH did not make significant changes to its staff management practices to accommodate this project, and similar efficiencies should be attainable at other institutions. Newly trained staff are not expected to achieve maximum speeds immediately, but it is our experience that staff begin achieving personal maximal efficiency after only 2-3 weeks of daily imaging, sometimes sooner.

Additionally, the described workflow does not include data entry, nor does it rely on the presence of previously captured data. Images captured by the photostation are transferred to HUH preservation storage nightly and ingested into the HUH specimen database to be used for data entry using a locally developed transcription web application. Quality control of images is performed daily by selecting a random sample of images from each batch. Images are inspected for deviations in focus, exposure, color, orientation, the presence of hands or fingers, and any other noticeable issues. After inspection, images are processed in batch with minor adjustments for white balance, exposure, levels, and lens correction. Derivative images are also created during this post-processing stage. Performing these steps takes the operator approximately five minutes per batch, while the creation of derivative images may take 30 minutes to two hours (unattended), depending on the available computing power and size of the batch.

After image post-processing is complete, images are passed into an informatics pipeline for additional transformations, data extractions, and ingest into the HUH database and transcription application. Barcodes are automatically read from the image by software during ingest and the image is linked if there is an existing database record. In cases where multiple specimens are affixed to the same sheet, the system will identify each barcode and link the image to all pre-existing records. Images linked to records at this time are immediately available through the HUH website. Other components of the pipeline under active development include automatic image border cropping, label identification, and deep learning handwriting recognition models.

## Key design and enhancement features to earlier imaging approaches

By decoupling the imaging and subsequent label transcription stages of the digitization workflow, which are often performed side-by-side, we were able to focus exclusively on fine-tuning the imaging workflow and apparatus, resulting in our reported efficiencies. While some herbarium workflows may partially, or mostly, decouple imaging from data capture (Heerlien et al., 2015; Thiers et al., 2016), the need for any data capture, even for skeletal record creation, can add time to the imaging workflow or require custom and sometimes poorly supported software additions. By fully decoupling image capture from data creation, relying instead on feeding images to the backend informatics pipeline, we have reduced workflow complexity for image capture and avoided the need for custom software on the photostation. Moreover, our workflow has greatly simplified the steps involved so there is no ambiguity as to where a specimen exists in the digitization workflow or what steps an operator should execute. Specimens are imaged first and immediately refiled in small batches in close proximity to the collection. This reduces the time specimens are kept out of the collection and, as indicated above, allow staff to focus solely on imaging without coordinating imaging and transcription in a single process. Efficiencies gained using this new photostation are further achieved via numerous ergonomic customizations specific to the handling and dimensions of standard herbarium sheets, the inclusion of components that have a high return on investment at scale (e.g., automatic barcode dispenser), and the use of automated image processing steps to replace manual activity (e.g., image recognition for barcode data entry). Key customizations include: platform sizing for vascular sheets to eliminate manual turning of specimens, adjustable shelving to easily separate in-process from completed specimens (queue tray versus receiving area), remote camera shuttering, height adjustment for operator comfort and efficiency, eye-height tablet display for quickly assessing image quality, and built-in automatic barcode dispenser. Additionally, the workflow streamlines image processing steps by moving these activities to a later stage where they can be performed in batch and with the aid of automation.

In addition to these enhancements, the shape and size of the photostation is a significant benefit of our design. Measuring only 189cm tall, 92cm wide, and 62cm deep and requiring only access to a power outlet, institutions adopting this design are likely to find a suitable space near their collection to deploy this hardware. At the HUH, a single digitization area requires only enough space to accommodate the photostation, operator, herbarium cart, and optional worktable (Fig. 1). The photostations deployed at the HUH also use a wired internet connection, but Wi-Fi can be used depending on the number of images captured daily and the speed of the wireless network for transferring the images to institutional storage. This photostation design also requires no complex installations or long-term space commitments. It can be moved or placed into storage in response to changing institutional priorities or available funding. Additional units can also be constructed, as at the HUH, to increase the overall rate of digitization or respond to new projects as they arise. Importantly, with this revised hardware and improved workflow we have dramatically increased the cost-effectiveness of imaging while eliminating the space and labor obstacles associated with the conveyor belt design, thus facilitating the deployment of lightweight, mobile photostations throughout the collection range.

## Comparison of imaging efficiency to other approaches

The imaging workflow and photostation summarized above has been in operation at the HUH since mid-2018 (first prototype deployed fall of 2017). This gives us many thousands of hours of staff time to analyze our imaging performance and to make clear comparisons in efficiencies gained relative to previously implemented imaging workflows, including those with more traditional single-user interfaces and the conveyor belt system described above. It should be mentioned that herbarium digitization workflows are diverse and challenging to summarize (Nelson et al., 2015). However, Thiers et al. (2016) and Nelson et al. (2012), who adopted commonly applied single-station approaches, reported average imaging rates of around 100 images (not including label data capture) per hour (or ca. 36 seconds per specimen) per imaging station. The conveyor belt workflow described by Sweeney et al. (2018) was tested for “image-only” throughput where only a barcode or QR code was scanned in addition to the image capture. These tests achieved an impressive 20 seconds per image (or ca. 180 specimens per hour), a substantial improvement over previous efforts using single user interfaces. In contrast, our refined workflow and single-user photostation captures images with comparable minimal data capture (a barcode) at an estimated average of 8 seconds per image. Requiring half the staff to operate, our average rate of image capture is a four-fold increase in efficiency over the best reported times of the conveyor-belt system.

## Broader implications of high-throughput imaging

Herbaria undertaking digitization projects must address difficult questions about project scope and focus, and importantly how to retrieve relevant specimens from the collection in a way that is efficient and cost-effective. Current funding sources for herbarium digitization, such as NSF ADBC, often require a thematic focus but many novel research themes are not congruent with typical models of herbarium organization, which are usually organized by family or geography. As outlined above, the HUH has participated in many such digitization projects but answering questions about dispersed subsets of the collection is often labor intensive. For instance, a list of counties spanning ten U.S. states was used to circumscribe the scope of the SoRo TCN. Sometimes, these questions cannot be answered based on the amount of labor and availability of staff, and accuracy is often questionable. Because of these necessary thematic circumscriptions, staff undertaking digitization must sort through numerous specimens to identify only those of relevance to the project, at a significant loss to overall efficiency.

At the HUH, the development of our photostation greatly propelled our effort to undertake a key mission outlined in the 2018 HUH Strategic Plan: digitizing the entirety of our North American vascular plant holdings. This represents ca. 1.5 million specimens in our collections including among the oldest and most extensive collections of North American plants in the world. To date, HUH staff have imaged and databased over 500,000 specimens using the described photostation and workflow. This ambitious goal has further benefited by funding from several completed and ongoing NSF-funded TCNs, including: the above mentioned NEVP, ‘Using Herbarium Data to Document Plant Niches in the High Peaks and High Plains of the Southern Rockies: Past, Present, and Future’ (SoRo), ‘Digitizing “endless forms”: Facilitating Research on Imperilled Plants with Extreme Morphologies’ (PoE), and ‘American Crossroads: Digitizing the Vascular Flora of the South-Central United States’ (TORCH). More immediately, our success with implementing this workflow organization was essential in remaining operational during recent COVID shutdowns, in which HUH staff were unable to return to campus for three months. In advance of the shutdown, HUH staff prioritized imaging activities and created a backlog of images that allowed approximately 13 staff to continue working full-time on transcription and georeferencing activities while remote. At the time of this writing (9/2020), the HUH is operating under a low-density occupancy plan. However, because of the speed of imaging, and as noted above, staff only need to return to the office once per week to generate enough specimen images for full-time data capture activities.

Our photostation has been a boon to productivity and to our current NSF-funded efforts and to the larger goal to digitize the entirety of our North American holdings. Our capacity to affordably image broad sections of the collection with minimal data capture has benefitted the HUH in numerous ways. In cases where we have already imaged a significant portion of a collection that is relevant to a project, we can generate precise counts by taxon or geography (county-level and above) to accurately estimate the labor required and funding necessary for further digitization (usually full-label transcription and georeferencing). Using our backend software, it is trivial to present staff with project-specific subsets of specimen images for full transcription and georeferencing, which are by far the most time-consuming stages of these workflows. By targeting entire families or continents for “initial” digitization, the HUH has been able to participate in many overlapping digitization efforts without managing multiple workflows.

We have also identified that our efforts have stimulated tremendous interest in these data, signaling that there is no shortage in the demand for herbarium specimen data. Between 2017, when the North America digitization project was launched, and 2019, the HUH saw a 260% increase in citations through GBIF. This is despite the fact that over 50% of these newly created ca. 500,000 records only include a taxon and state (or county). We believe that by advancing the development of extremely cost-effective solutions to herbarium specimen digitization, even with minimal data capture, the community can increase their utility and service to scientific advancement, far beyond what we do today.

Along these lines, our photostation design and workflow are notable for their greatly increased efficiency without reliance on a striking technical break-through. The assembly of our photostation, while novel in its refinements, uses commonly available hardware and electronics. The informatics pipeline and backend, briefly noted here, perform similar functions to other existing workflows. Indeed, while these refinements have yielded increases in efficiency, much has been gained in the organization and optimization of our workflow achieved via our iterative design and prototyping process. We hope this effort will allow others to benefit from our workflow. For example, institutions with smaller budgets or seeking to leverage existing photographic equipment could deploy a light table in a room with color-matched lighting to reproduce some of the efficiencies gained by quickly staging and de-staging the specimen. Accessories like a quick-fire USB button or automatic barcode dispenser could also be deployed within reach to save seconds on each image capture. Likewise, data capture can occur after bulk imaging activities if the software tools are available. In fact, this ad-hoc configuration is similar to what we used to validate our initial hypothesis about possible efficiency gains.

## Conclusions

The inability to quickly and cost-effectively image herbarium specimens has been an impediment to the broader digitization of the world’s herbaria. Herbaria have made tremendous strides in mobilizing these data in the past decade, which has been greatly facilitated by community involvement and broad participation. We feel this photostation marks a tremendous improvement to current efforts. It can be easily scaled to larger operations by producing more individual stations or reduced accordingly for smaller projects. In addition, the return to a single user interface obviates the need for conveyor belt apparatuses which are often expensive (especially when a vended solution is weighed), space consuming, and especially reliant on a constant work staff. All of these factors are liable to be substantial barriers to entry for most institutions. We feel that this advancement is particularly relevant and timely because there exist more than 390 million herbarium specimens in the world and only a small fraction of these specimens have been imaged to date (Sweeney et al., 2018). Here, we freely distribute the design and plans of this apparatus to expand efforts to more easily and efficiently liberate collections that have been hidden from view for decades to centuries.

## Author contributions

CCD initiated, supervised, and worked to fund the effort to expand specimen digitization during his tenure as HUH Director (2013-2019) and as Curator of Vascular Plants (ongoing). JK and CJG conceived the original idea and high-level design for the photostation hardware and imaging workflow with significant intellectual input from CCD and HUH staff. CJG designed and constructed the first photostation prototype with input and improvements by JK. JK supervised the day-to-day management of the project (ongoing), researched imaging technologies and configurations, developed the software components, and performed the data analysis. CCD and JK wrote the manuscript with final acceptance by all authors.

## Acknowledgements

We acknowledge Michaela Schmull, Anne Marie Countie, and the HUH curatorial staff who greatly helped to facilitate our design and implementation of the photostation. Special thanks to Anthony Brach, Ellie Taylor, and Emma Tanner who provided important details on the photostation workflow. This work was largely conducted on the traditional territory of the Wampanoag and Massachusetts people. Funding was provided by the HUH, and by the ADBC program of the U.S. National Science Foundation (awards 1208835 [NEVP], 1702322 [SoRo], 1802209 [PoE], and 1902078 [TORCH]). We are indebted to L. Caria for his assistance with Figure 1.

## Notes

### Competing Interest Statement

The authors have declared no competing interest.

